# The LINC complex and microtubule motors regulate the number and position of nuclei in the subperineurial glial cells of the *Drosophila* blood-brain barrier

**DOI:** 10.1101/2025.10.29.685339

**Authors:** Olivia R. Annes, Anton Schmitt, Daniel B. Akinremi, Daniel Koskas, Yunshu Qiu, Hanna Jewell, Jeffrey M. DaCosta, Eric S. Folker

## Abstract

Multinucleated cells, or syncytia, provide a unique system in which to understand the mechanisms of cellular organization. The two most dramatic features of syncytial cells are the number of nuclei and the positioning of nuclei within a shared cytoplasm. While the mechanisms that regulate these features have been studied in some syncytial cells, most syncytial cells are uncharacterized. Furthermore, whether the formation of the syncytia and the organization of the syncytia are linked is not known. We have characterized the subperineurial glial cells (SPG) which form the most restrictive layer of the *Drosophila* blood-brain barrier. We have found that disruption of the Linker of Nucleoskeleton and Cytoskeleton (LINC) complex, Kinesin, or cytoplasmic Dynein specifically in SPG cells affected both SPG cell development and general brain development. Specifically, the brains were smaller in each case and the SPG cells were smaller when the LINC complex or cytoplasmic Dynein were disrupted. The number of nuclei per cell was increased when Kinesin was disrupted, decreased when cytoplasmic Dynein was disrupted, and abnormal numbers of nuclei were found when the LINC complex was disrupted. Finally, the positions of nuclei relative to their nearest neighbor was decreased when the expression of each gene was disrupted and nuclei were closer to the cell edge when either Kinesin or cytoplasmic Dynein were disrupted. Finally, the evenness of nuclear spacing was reduced when LINC complex or Kinesin expression was disrupted. Together, these data illustrate that formation of SPG cells and the organization of SPG cells are dependent on microtubule motors and the LINC complex.

## Introduction

A fundamental goal of cell biology is to define the structural features of cells and the molecular mechanisms that give rise to these emergent features. Historically, the cell nucleus has been depicted as a spherical organelle housing the genome near the cell center. However, the mechanisms that drive dynamic changes in the position of the nucleus have emerged as critical to disparate cellular processes including cell division (Tran *et al*., 2001; Hazelwood *et al*., 2025), cell migration (Gomes *et al*., 2005), and cell differentiation (Petkova *et al*., 2019) among others (Gundersen and Worman, 2013; Bone and Starr, 2016). Mechanistically, the movement of nuclei is dependent on the Linker of Nucleoskeleton and Cytoskeleton (LINC) complex in most cells (Lee and Burke, 2018). The LINC complex is composed of Sad1, Unc84 (SUN) domain proteins that span the inner nuclear membrane, and Klarsicht, Anc, and Syne Homology (KASH) domain proteins that can interact with SUN domain proteins in the perinuclear space, thus concentrating much of their localization to the outer nuclear membrane (McGee *et al*., 2006; Sosa *et al*., 2012; Hao *et al*., 2021). These proteins enable mechanotransduction between the nucleoplasm and the cytoplasm. Within the context of nuclear movement and position the LINC complex harnesses mechanical work produced by the cytoskeleton to move the nucleus (Brosig *et al*., 2010; Janota *et al*., 2020). In many cell types, the force to move the nucleus is provided by the microtubule cytoskeleton (Tran *et al*., 2001; Tsai *et al*., 2010; Grava *et al*., 2011; Cadot *et al*., 2012; Metzger *et al*., 2012; Wilson and Holzbaur, 2012; Collins *et al*., 2021).

Although the mechanisms that regulate the position of the nucleus have been the subject of much research in recent years, the cell types that have been studied are relatively few, leaving it possible that different cell types have evolved unique mechanisms to suit their specific needs. This idea is well-illustrated by comparing migrating fibroblasts that utilize actin retrograde flow to move their nucleus (Gomes *et al*., 2005; Luxton *et al*., 2010) to migrating cortical neurons that utilize microtubule motors (Tsai *et al*., 2010). Thus, the mechanisms of nuclear movement and positioning must be tested in each cell type.

Multinucleated cells, or syncytia, provide dramatic systems in which to study nuclear movement. These cells exist across the eukaryotic lineage ranging from filamentous fungi (Mela *et al*., 2020), developing blastoderms (Foe and Alberts, 1983), differentiated myofibers (Abmayr and Pavlath, 2012), osteoclasts (Sabe *et al*., 2024), and syncytiotrophoblasts (Renaud and Jeyarajah, 2022), and pathogenic responses to cancer (Ogawa *et al*., 2025) and viral infection (Leroy *et al*., 2020). Because multiple nuclei share a single cytoplasm in these cells, the position of each nucleus cannot be defined only by its relationship to the cell center as is commonly done in mononucleated cells. The position of nuclei relative to one another, the cell borders, and other cell specific hallmarks also define the position of each nucleus and the overall nuclear spacing within the cell.

Few syncytial cells have been investigated with regard to the mechanisms that regulate their nuclear spacing (Padilla *et al*., 2022). Two relevant exceptions are differentiated myofibers and developing blastoderms. In addition to the LINC complex, the spacing of nuclei in myofibers is dependent on microtubules, microtubule motors, various microtubule associated proteins, and in mammalian myofibers the intermediate filament protein Desmin (Cadot *et al*., 2012; Elhanany-Tamir *et al*., 2012; Folker *et al*., 2012, 2013; Metzger *et al*., 2012; Wilson and Holzbaur, 2012, 2014; Falcone *et al*., 2014; Collins *et al*., 2017, 2021; Roman *et al*., 2017; Mandigo *et al*., 2019). Similarly, the spacing of nuclei in the syncytial blastoderm is dependent on microtubules, a different set of microtubule motors, and a different set of microtubule associated proteins (Foe and Alberts, 1983; Callaini *et al*., 1992; Deneke *et al*., 2019; de-Carvalho *et al*., 2022). Whether either of the previously described mechanisms apply to other syncytial cells has not been investigated.

The subperineural glial (SPG) cells of the *Drosophila* blood-brain barrier (BBB) present another syncytial system for studying the mechanisms of nuclear spacing. These cells, which compose the septate junction forming layer of the BBB, are flat and significantly larger than other glial cells (Stork *et al*., 2008; Von Stetina *et al*., 2018). While the SPG cells in the ventral nerve cord are mononucleated, the SPG cells that line the optic lobe undergo endomitosis throughout larval development, thus creating syncytial cells that support the necessary increase in total cell area without cell division and the disruption of the septate junction borders (Babatz *et al*., 2018; Von Stetina *et al*., 2018).

To gain a greater insight toward the mechanisms that regulate nuclear spacing in syncytia and how these mechanisms relate to cellular development we have screened several different genes that we previously demonstrated as crucial for nuclear spacing in *Drosophila* myofibers to determine whether they are also necessary to space nuclei in subperineurial glial cells in the *Drosophila* blood-brain barrier. We find that although nuclear spacing is LINC complex- and kinesin-dependent in both cell types, SPG cells do not require Ensconsin/MAP7 indicating a critical difference in the development and organization of these two syncytial cell types.

## Results and Discussion

The larval *Drosophila* brain is composed of the ventral nerve cord (VNC) and the spherical optic lobes (Figure 1A). Whereas the SPG cells that line the VNC were previously shown to be mononucleated, the SPG cells that line the optic lobes can be either mononucleated or multinucleated (Von Stetina *et al*., 2018). Because the internal organization of these cells has not been investigated, we first measured the distribution of nuclei in multinucleated SPG cells that line the optic lobes. Furthermore, our subsequent experiments utilize Moody-GAL4 to drive SPG-specific expression of RNAi constructs to test for cell-autonomous effects. Therefore, we used Moody-GAL4 driven expression of UAS-mCherry RNAi as a control and measured the effects on viability, brain size, SPG cell size, and nuclear position in SPG cells.

**Figure 1.**
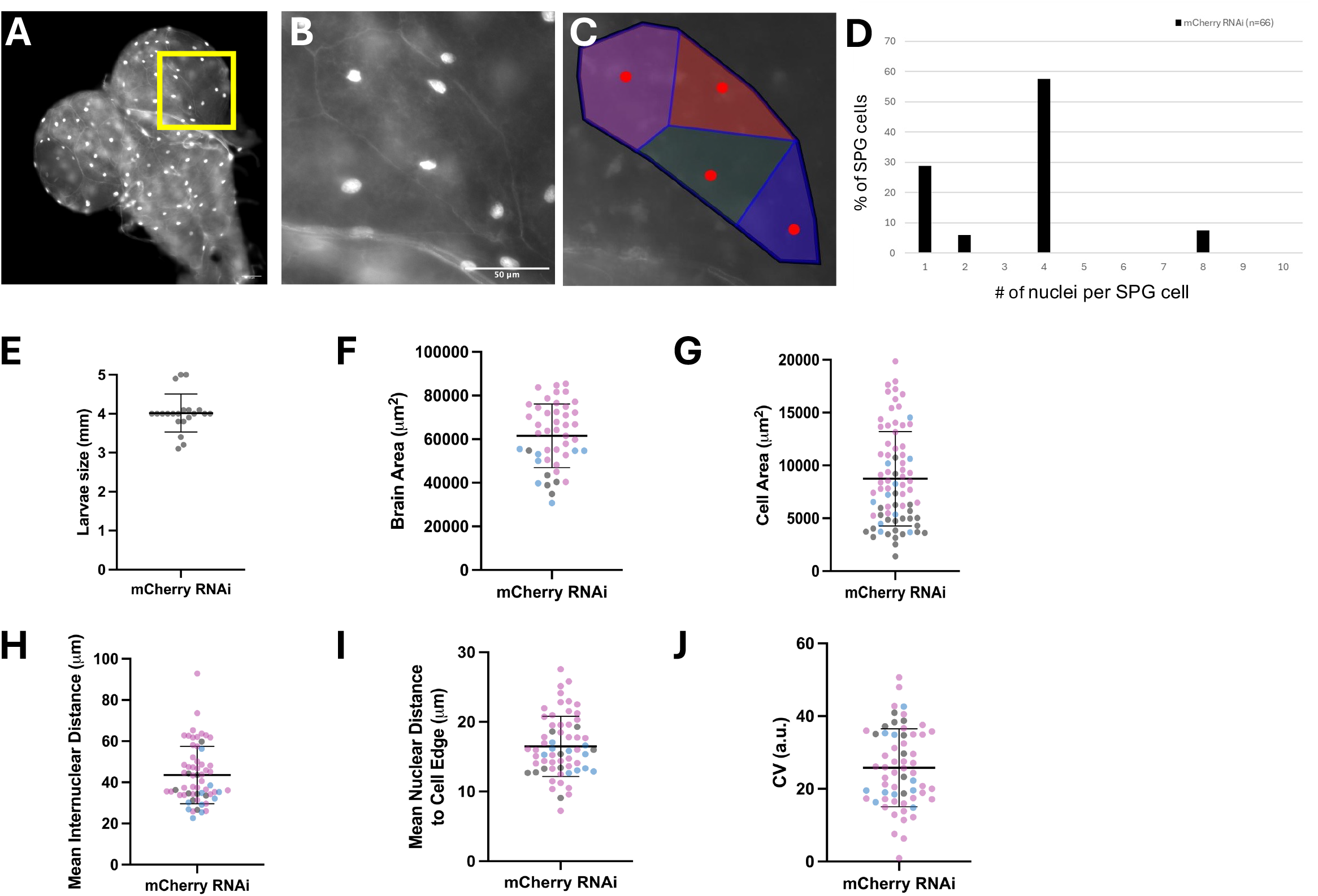
Characterization of nuclear position in SPG cells. A. Full brain of L3 larva expressing UAS-NLS-GFP under the control of Moody-GAL4 to identify nuclei of the SPG cells and NrxIV-GFP to identify the septate junctions that are specific to the SPG cell-cell contacts. B. Single SPG cell from yellow boxed region in A. C. Voronoi polygons overlayed on the single SPG cell shown in B. All scale bars are 50 µm. D. Histogram indicating the number of nuclei per SPG cell. Data indicate the percent of total cells counted (n=77). E. Length of L3 larvae. Data points represent single larvae (n=21). F. Cross-sectional area of optic lobes of control brains. Data points represent one optical lobe (n=45). G. Cross-sectional area of individual SPG cells. Data points represent a single SPG cell (n=77). H. Mean distance from each nucleus to its nearest neighbor. Data points represent the average value for all nuclei within a single cell (n=61). I. Mean distance from each nucleus to the nearest cell-cell contact. Data points represent the average value for all nuclei within a single cell (n=61). F-J. Different colors represent different biological replicates.

Animals that expressed mCherry RNAi in SPG cells were an average of 4 mm long with a range of 3 mm to 5 mm (Figure 1E) and the optic lobes had an average cross-sectional area of 60,000 µm^2^ with a range of between 40,000 µm^2^ and 80,000 µm^2^ (Figure 1A,F). The areas of the individual SPG cells were more variable, with an average of approximately 9,000 µm^2^ but a range of between 1,000 µm^2^ and 20,000 µm^2^ (Figure 1B,G). This variation in size correlated with the number of nuclei, as larger cells tended to have more nuclei (Figure 2I, 3I).

**Figure 2.**
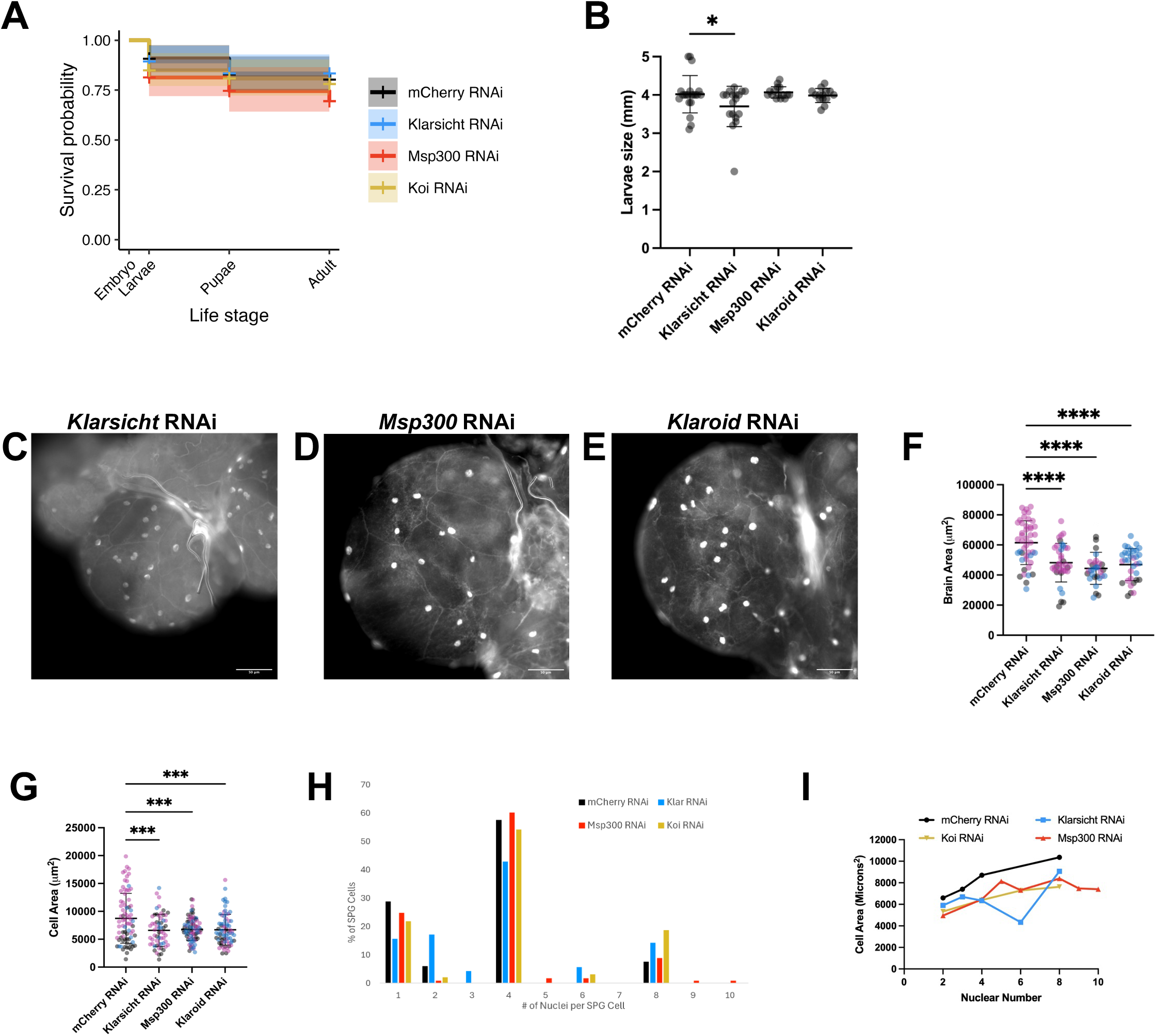
LINC complex expression in SPG cells regulates brain development. A. Survival probability of indicated genotypes. Lines indicate mean from 5 different experiments and shading indicates 95% confidence intervals. B. Length of L3 larvae in millimeters. Data points represent a single larva. C-E. Fluorescence images of the nuclei and septate junctions in the SPG cells from an optical lobe of the brain from an L3 larva that expressed RNAi against Klarsicht (C), Msp300 (D), and Klaroid (E). All scale bars are 50 µm. F. Cross-sectional area of brain optic lobes. Data points represent one brain lobe. G. Cross-sectional area of SPG cells. Data points represent a single SPG cell. H. Histogram of the number of nuclei per cell. Bars represent % of SPG cells, mCherry RNAi n=66, Klar RNAi n=70, Msp300 RNAi n=113, Koi RNAi n=96. I. Graph of cell area vs nuclear number. Data points represent the mean area for all SPG cells with the indicated number of nuclei. F/G. Different colors indicate different biological replicates.

To define the positions of nuclei within the individual SPG cells we made several measurements. First, we measured the distance between each nucleus and its nearest neighbor nucleus which had a mean of 42 µm and a range of 21 µm – 95 µm (Figure 1H). Similarly, we measured the distance from each nucleus to the nearest cell edge and found that nuclei were on average 18 µm from the edge with a range of 8 µm – 28 µm from the cell edge (Figure 1I). Finally, to measure how evenly distributed nuclei were, we created Voronoi polygons in which each pixel in a cell is assigned to the nearest nucleus, indicated by differently colored polygons surrounding each nucleus (Figure 1C). We then calculated the coefficient of variation (CV) between the areas of the polygons ascribed to each nucleus (Figure 1J). The mean CV was approximately 25 with a range of 0 to 55.

Our finding that the nuclei are positioned similarly in SPG cells from dozens of brains across multiple biological replicates indicated that the positions of nuclei are actively regulated in SPG cells. We next aimed to identify the genes that regulate this spacing. Our first focus was the LINC complex which regulates nuclear position in diverse systems (Lee and Burke, 2018). Furthermore, we and others have shown that the LINC complex components *Msp300, Klarsicht*, and *Klaroid* all regulate nuclear spacing in *Drosophila* syncytial myofibers (Elhanany-Tamir *et al*., 2012; Mandigo *et al*., 2019; Padilla *et al*., 2025). We expressed RNAi against two genes encoding KASH domain proteins, *Msp300* and *Klarsicht*, and one gene encoding a SUN domain protein, *Klaroid*. We have validated each of these RNAi lines previously (Mandigo *et al*., 2019; Padilla *et al*., 2025). We first assessed the effects of each RNAi on *Drosophila* viability, brain morphology, and SPG cell size (Figure 2). Viability was not affected when any of the LINC complex components were depleted from SPG cells (Figure 2A). We did note a small change in the average size of animals that expressed RNAi against *Klarsicht*, but this is based on a single outlier that was much smaller than any other animal measured (Figure 2B). We therefore concluded that the expression of the LINC complex in SPG cells does not regulate larval size. Conversely, the size of the optic lobes was significantly reduced when RNAi was expressed against any of the 3 LINC complex components (Figure 2C-F). Consistent with this, the average size of individual SPG cells was also reduced when the expression of LINC complex encoding genes was disrupted (Figure 2G).

To determine whether the reduced SPG cell size correlated with a reduction in the number of nuclei, we counted the number of nuclei per SPG cell. Disruption of *Klarsicht* expression increased the number of SPG cells with 2 nuclei and decreased the number of SPG cells with 4 nuclei (Figure 2H). Conversely, the disruption of *Klaroid* expression increased the number of SPG cells with 8 nuclei and the disruption of *Msp300* expression had almost no effect on the number of nuclei per SPG cell (Figure 2H), suggesting that cell size and brain size are not strictly tied to the number of nuclei in each SPG cell when LINC complex expression is disrupted. Altogether, these data indicate that while viability and the developmental timeline are not affected by the loss of the LINC complex from SPG cells, the development of the SPG cells themselves, and the brain more generally, requires both KASH domain proteins (Klarsicht and Msp300) and the SUN domain protein Klaroid.

We next examined the internal organization of multinucleated SPG cells that expressed RNAi against each LINC complex component (Figure 3A-C). The distance between neighboring nuclei was reduced when the expression of any of the three LINC complex components was disrupted (Figure 3D). Conversely, the distance between each nucleus to the nearest cell edge was not affected in cells where the expression of the LINC complex was disrupted (Figure 3E). Finally, to determine how evenly nuclei were distributed in SPG cells me measured the CV of the Voronoi polygons associated with each nucleus (Figure 3A’-C’). The CV was increased when the expression of any of the three LINC complex genes was disrupted. Together, these data indicate that the LINC complex is a critical regulator of nuclear spacing in SPG cells. More specifically, the LINC complex regulates the distance between nuclei, but not the distance between the nucleus and the nearest cell-cell contact.

**Figure 3.**
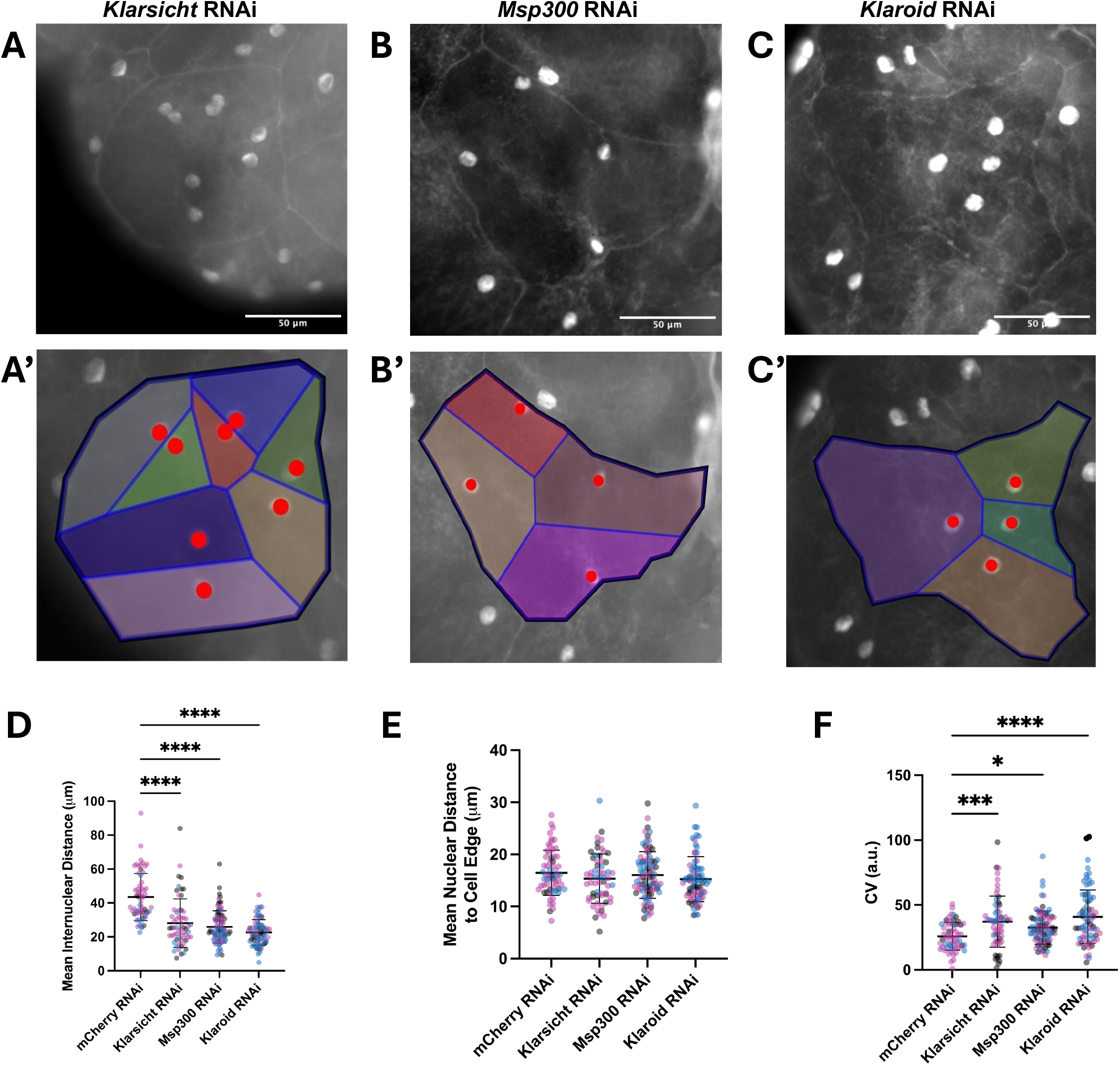
LINC complex expression in SPG cells regulates the number and position of nuclei A-C. Fluorescence images of the nuclei and septate junctions in individual SPG cells (A-C) and the same images with Voronoi polygons overlaid (A’-C’) in brains from animals that expressed RNAi against Klarsicht (A), Msp300 (B) or Klaroid (C). D. Mean nuclear distance from each nucleus to the nearest nucleus. Data points represent the average distances of all nuclei within a single cell. E. Mean nuclear distance from each nucleus to the nearest cell-cell contact. Data points represent the average distances of all nuclei within a single cell. F. Coefficient of variation in Voronoi polygon size. Data points represent a single SPG cell. D-F. Different colors represent different biological replicates.

Nuclear positioning in many cells, including syncytial muscle cells and syncytial blastoderms is also dependent on microtubules and microtubule motors (Gundersen and Worman, 2013; Padilla *et al*., 2022). Therefore, we hypothesized that nuclear position in SPG cells would also be microtubule-, Kinesin-, and Dynein-dependent. To test this hypothesis, we expressed RNAi against three genes critical for nuclear positioning in *Drosophila* myofibers: Kinesin heavy chain (*Khc*), Dynein heavy chain 64C (*Dhc64C*), and Ensconsin (*Ens*). *Khc* and *Dhc64C* encode the microtubule motors Kinesin-1 and cytoplasmic Dynein, respectively, whereas *Ens* encodes *Drosophila* MAP7.

Expression of RNAi against *Dhc64C* greatly reduced viability during the larval stage of development (Figure 4A). Furthermore, the animals that did survive were much smaller than controls or animals that expressed any other RNAi (Figure 4B). Similarly, when *Khc* expression was disrupted in SPG cells, *Drosophila* viability was reduced (Figure 4A) and the surviving animals were smaller compared to controls (Figure 4B) indicating that both microtubule motors are critical for SPG cell function. However, the disruption of *Ens* expression did not affect viability or growth (Figure 4A,B). Together these data suggest that cytoplasmic Dynein and Kinesin expression in SPG cells is critical for viability and growth.

**Figure 4.**
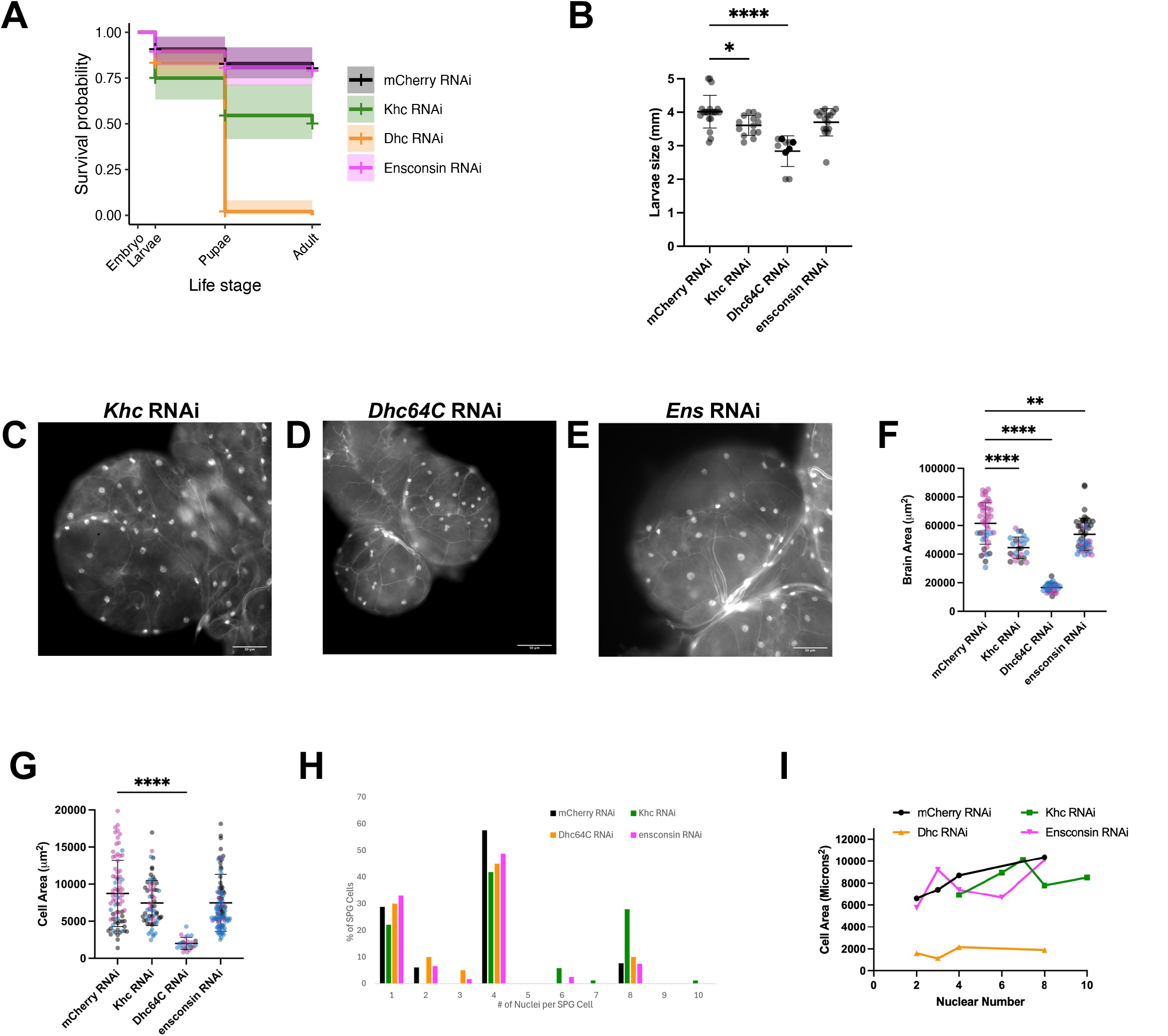
Microtubule motors in SPG cells regulate brain development. A. Survival probability of indicated genotypes. Lines indicate mean from 5 different experiments and shading indicates 95% confidence intervals. B. Length of L3 larvae in millimeters. Data points represent a single larva. C-E. Fluorescence images of the nuclei and septate junctions in the SPG cells from an optical lobe of the brain from an L3 larva that expressed RNAi against Khc (C), Dhc64C (D), and Ens (E). All scale bars are 50 µm. F. Cross-sectional area of brain optic lobes. Data points represent one brain lobe. G. Cross-sectional area of SPG cells. Data points represent a single SPG cell. H. Histogram of the number of nuclei per cell. Bars represent % of SPG cells, mCherry RNAi n=66, Khc RNAi n=69, Dhc64C RNAi n=20, Ens RNAi n=121. I. Graph of cell area vs nuclear number. Data points represent the mean area for all SPG cells with the indicated number of nuclei. F/G. Different colors indicate different biological replicates.

We next examined the brains specifically. The optic lobes of the brain were significantly smaller than controls or animals that expressed RNAi against any of *Dhc64C, Khc*, or *Ens* (Figure 4C-F). Similarly, the individual SPG cells were also smaller when the expression of *Dhc64C, Khc*, or *Ens* were disrupted, but the difference was only statistically significant when *Dhc64C* expression was disrupted (Figure 4G). Consistent with the general developmental effects, disruption of expression of *Khc, Dhc64C*, or *Ens* affected the number of nuclei per SPG cell (Figure 4H). In controls, nearly 60% of SPG cells contain 4 nuclei. Another 20% of control SPG cells contain only 1 nucleus, with most of the mononucleated cells being near the VNC as previously described (Von Stetina *et al*., 2018). The remaining 10% of control SPG cells contain either 2 nuclei or 8 nuclei. Disruption of *Khc* expression caused an increase in the percentage of cells with 8 nuclei whereas disruption of *Dhc64C* expression caused an increase in the percentage of cells with 2 nuclei. These data suggest that reduced *Khc* expression increased the number of mitotic cycles whereas reduced *Dhc64C* expression decreased the number of mitotic cycles. More interestingly, expression of RNAi against any of *Khc, Dhc64C*, or *Ens* resulted in cells with abnormal numbers of nuclei. Whereas SPG cells in controls always had 1, 2, 4, or 8 nuclei, cells with 3 nuclei, 6 nuclei, 7 nuclei, or 10 nuclei were found when any of the RNAi constructs were expressed (Figure 4H). Furthermore, the scaling between cell size and number of nuclei (Figure 4I) was dramatically affected by the expression of RNAi against *Dhc64C*, possibly explaining the significant effects on viability. These data indicate that the microtubule cytoskeleton, and microtubule motors more specifically regulate the number of nuclei in SPG cells.

We next investigated whether *Khc, Dhc64C*, and *Ens* regulate nuclear positioning in SPG cells (Figure 5 A-C). Expression of RNAi against either *Khc* or *Dhc64C* caused a decrease in the internuclear distance (Figure 5D) and distance from nucleus to cell edge (Figure 5E). However, only the disruption of *Khc* expression caused a statistically significant increase in the CV of the Voronoi polygon areas associated with each nucleus (Figure 5F). Because the CV of the Voronoi polygon areas controls for changes in cell size, we conclude that *Khc*, but neither *Dhc64C* nor *Ens*, regulates the spacing between nuclei in SPG cells. However, *Dhc64C* does regulate how far nuclei are from the cell borders.

**Figure 5.**
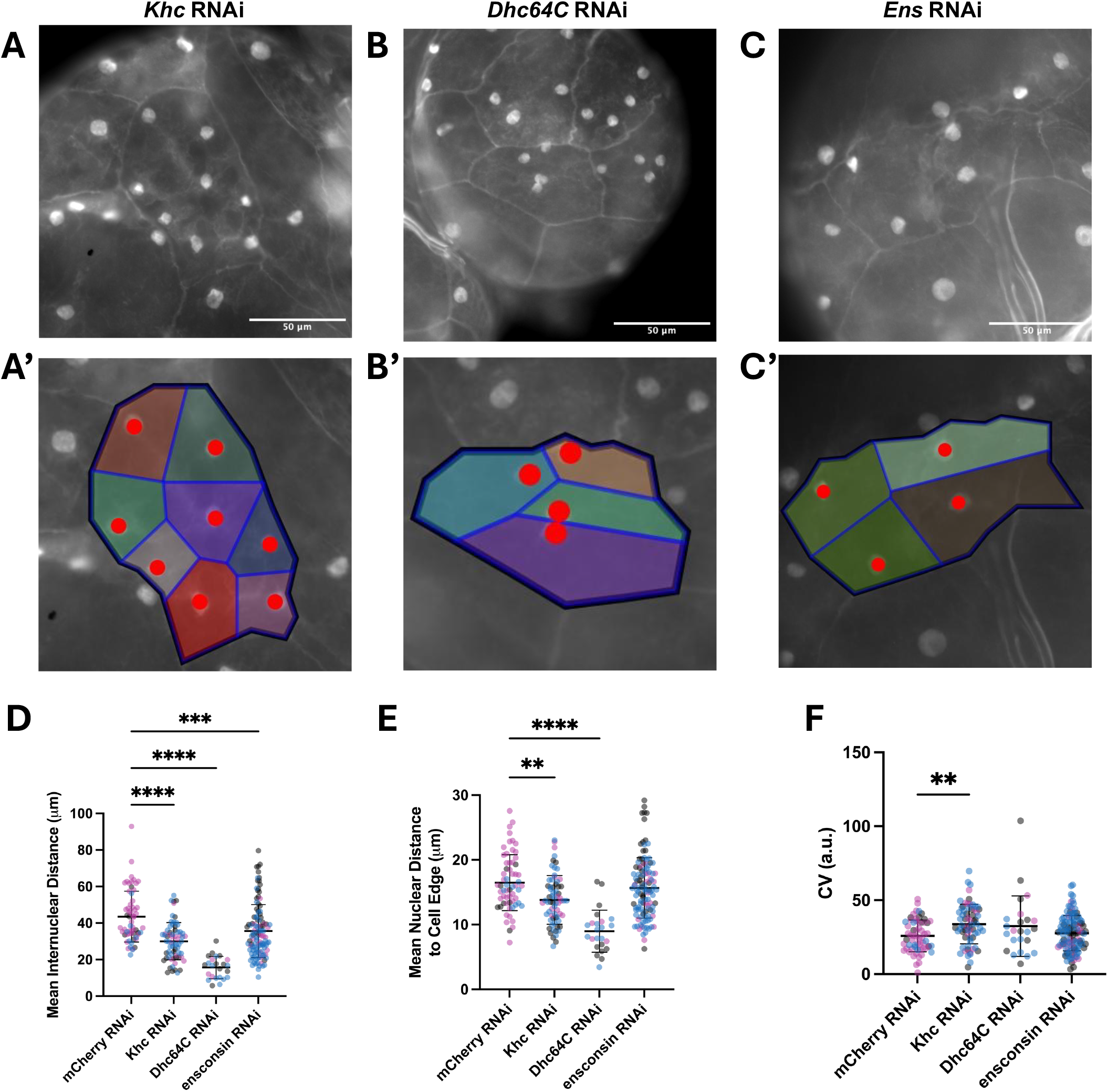
Microtubule motors in SPG cells regulate the number and position of nuclei A-C. Fluorescence images of the nuclei and septate junctions in individual SPG cells (A-C) and the same images with Voronoi polygons overlaid (A’-C’) in brains from animals that expressed RNAi against Khc (A), Dhc64C (B) or Ens (C). D. Mean nuclear distance from each nucleus to the nearest nucleus. Data points represent the average distances of all nuclei within a single cell. E. Mean nuclear distance from each nucleus to the nearest cell-cell contact. Data points represent the average distances of all nuclei within a single cell. F. Coefficient of variation in Voronoi polygon size. Data points represent a single SPG cell. D-F. Different colors represent different biological replicates.

Many large cell types are syncytial, and the mechanism of multinucleation has been described for many systems. For example, myofibers and syncytiotrophoblasts form from the fusion of mononucleated precursor cells (Abmayr and Pavlath, 2012; Renaud and Jeyarajah, 2022). Conversely, SPG cells and developing insect blastoderms form from endomitosis (Foe and Alberts, 1983; Von Stetina *et al*., 2018). What is significantly less understood is whether the spacing of nuclei is regulated in each cell type, whether the mechanisms of nuclear spacing are conserved, and whether the mechanism of multinucleation impacts the mechanism of cellular organization.

Here we have demonstrated that, like myofibers, the nuclei in SPG cells are actively spaced, and we have measured this spacing in three different ways. First, we measured the distance between nuclei; second, we measured the distance from each nucleus to the nearest cell border; third we measured the evenness of their spacing using the CV of their Voronoi polygons.

Using these measurements and SPG-specific RNAi expression, we have identified several genes that are critical for nuclear spacing, both relative to the other nuclei and relative to the cell-cell contacts in SPG cells. Because this paper represents the first examination of the internal organization of SPG cells, we focused specifically on the spacing of nuclei and tested only candidate genes that were previously shown to regulate nuclear spacing in *Drosophila* myofibers. This approach enabled the direct comparison of two cell types that form syncytia via distinct mechanisms. We find that these disparate cells require overlapping but distinct sets of genes to space their nuclei. Like myofibers, SPG cells rely on the LINC complex and the microtubule cytoskeleton to space their nuclei. However, expression of RNAi against *Ens* in myofibers results in a striking clustering phenotype (Metzger *et al*., 2012) whereas the expression of the same RNAi in SPG cells had only a subtle effect. These data together indicate that the general mechanisms of microtubule and microtubule motor dependent positioning of nuclei are likely differentially regulated based on the unique features of different syncytia.

Perhaps not surprisingly, different genes were required to regulate distinct features of the nuclear spacing, similar to previous work in myofibers (Perillo and Folker, 2018). The relative importance of each aspect of nuclear spacing remains to be determined. However, in the syncytial blastoderm, a syncytia that forms via endomitosis like SPG cells do, it is the spacing of nuclei relative to one another, and more precisely the evenness of that spacing, that is regulated (de-Carvalho *et al*., 2022). In looking specifically at the CV data as a proxy for how evenly spaced nuclei are, we see that neither *Dhc64C* nor *Ens* were necessary.

One surprising outcome of these experiments was the variation in the numbers of nuclei in each SPG cell. Expression of RNAi against each of the genes tested, those encoding the LINC complex, microtubule motors, or microtubule associated proteins, shifted the distribution in the number of nuclei. While the shift toward increased numbers of nuclei when we expressed RNAi targetting *Khc* or *Klaroid* likely represent increased mitotic cycles, the reduced numbers of nuclei when RNAi was expressed against *Dhc64C* are consistent with reduced mitotic cycles. Surprisingly, RNAi against *Klarsicht* caused an increase in cells with both more nuclei and fewer nuclei compared to controls indicating a more general dysregulation of mitosis. Perhaps more intriguing is the presence of cells with 3, 5, 6, 7, 9, and 10 nuclei. Although time-lapse imaging of these cells has not yet been possible, the fact that in controls all cells have 1, 2, 4, or 8 nuclei suggests that mitotic cycles are synchronized. The finding of different numbers of nuclei in each of our experiments indicates that each of these proteins is crucial for synchronization. Whether that function is related to the effects on nuclear spacing remains to be determined.

Altogether these data establish a baseline for the investigation of cellular organization in general, and nuclear spacing specifically, in SPG cells of the *Drosophila* blood-brain barrier. This endomitosis-dependent cell type can serve as a model of differentiated syncytium, and can be compared to the fusion-dependent myofibers to continue building a comprehensive understanding of syncytial cells.

## Acknowledgements

We thank the Vienna Drosophila Resource Center (VDRC) at Vienna BioCenter Core Facilities (VBCF), member of the Vienna BioCenter (VBC), Austria and we thank the Bloomington Drosophila Stock Center for the materials used in this work. Thank you also to Bret Judson and the Boston College Imaging Core for Infrastructure and support.

## Materials and Methods

### *Drosophila* genetics

All stocks were grown under standard conditions at 25°C. Moody-GAL4, UAS-NLS-GFP; NrxIV-GFP (Von Stetina *et al*., 2018) were used in all experiments to both drive UAS-RNAi constructs and to visualize the nuclei and the septate junctions in SPG cells. UAS-RNAi lines used were mCherry (BDSC 35785), Msp300 (BDSC 32377), Klar (BDSC 28313), Ens (VDRC 106270), Koi (VDRC 108236), Khc (BDSC 35409), and Dhc64C (BDSC 36698). mCherry RNAi (BDSC 35785) was expressed as a control in all RNAi experiments.

### Drosophila viability assay

Embryos were collected on grape juice agar plates. Embryos were bleached and rinsed with water before stage 16 embryos were selected based on the appearance of the trachea as previously described (Metzger *et al*., 2012). L1 larvae that hatched were moved to a vial of standard BDSC cornmeal food. Vials were stored at 25 ºC for 3 days. L3 larvae were floated from the food by submersion in a solution of 10% sucrose. Additional vials from which L3 larvae were not harvested were used to count the number of larvae that developed into pupae and adults.

### Sample preparation and fluorescence microscopy

L3 larvae were collected as described in the *Drosophila* viability assay. Larvae were measured using a standard ruler with mm markings and then submerged in cold PBS and their brains were removed as previously described (Von Stetina *et al*., 2018). Isolated brains were fixed in 10% formalin for 15 minutes at room temperature, then washed and rinsed three times with PBS. Brains were mounted immediately after dissection. Samples were mounted between two glass coverslips using ProLong Gold Antifade Mountant (Life Technologies; P36930). Samples were left to cure for 24 hours before imaging.

Samples were imaged on Zeiss Axioimager Z2 using a 20X/0.8NA Plan APOCHROMAT objective and FL Filter Set 38 HE GFP shift free. After each brain was imaged, the coverslips were flipped over so that the other side of the brain could be imaged.

### Generation of Voronoi diagrams of nuclear positioning

CZIs were uploaded to FIJI (ImageJ) for analysis. Brain and cell border measurements were measured using the polygon tool. Nuclei were identified manually using the multipoint tool to indicate the center of each nucleus. Measurements were uploaded in a csv file to a Python program developed on the Google Colab platform. The program calculates and organizes final results including cell area, number of nuclei, internuclear distance, and nuclear distance to edge onto an Excel sheet. The program also performs Voronoi polygon analysis by using each nucleus centroid as a Voronoi seed and the cell borders as bounding constraints. For SPG cells with three or more nuclei, the Voronoi and voronoi_plot_2d functions from the scipy.spatial module are used to generate the Voronoi diagrams. For SPG cells with only two nuclei, a modified Voronoi constructing function is used to identify the midpoint between two nuclei and draw a perpendicular bisector through this midpoint, thereby dividing the cell into two Voronoi polygons. The areas of each generated Voronoi polygon are then used to calculate the coefficient of variation, which is also organized onto the same Excel sheet for further statistical analysis.

### Standard Measurements

Fluorescence images of dissected brains from L3 larvae were analyzed using Fiji software.

*Brain Size*: Optical lobe size was measured by drawing an ROI around the entire optical lobe and measuring the enclosed area.

*Cell Size*: SPG cell size was measured by drawing an ROI around an individual SPG cell identified by the NrxIV-GFP (septate junction) signal and measuring the enclosed area. *Internuclear Distance:* The centroids of nuclei were identified as part of the generation of Voronoi polygons. The distances between each centroid and the nearest centroid were then taken as the distance from a nucleus to its nearest neighbor. A separate distance was determined for each nucleus and the average distances within a cell was used to determine the internuclear distance measurement for each cell.

*Distance to edge:* The cell perimeter and the nucleus centroid positions determined as part of the Voronoi polygon measurements were used to identify the shortest distance between each nucleus and the cell edge. A separate measurement was made for each nucleus and the average distance within a cell was used to determine the distance to edge value for each cell.

### Statistics

The survival analyses were completed in R v4, using the packages survival and survminer to estimate and plot Kaplan-Meier curves and their 95% confidence intervals. All other statistics were completed using Prism 10.6.0. All comparisons were ANOVA with Tukey HSD post-hoc test.

